# Unveiling Gene Modules at Atlas Scale through Hierarchical Clustering of Single-Cell Data

**DOI:** 10.1101/2025.03.12.642774

**Authors:** Feng Tang, Zhongmin Zhang, Weige Zhou, Guangpeng Li, Luyi Tian

## Abstract

A major challenge in scRNAseq analysis is how to recover the biologically meaningful cell ontology tree and conserved gene modules across datasets. Data integration and batch-effect correction have been the key to effectively analyze multiple datasets, but often fail to disentangle cell states in heterogeneous samples, such as in cancer and the immune system. Here we present super single cell clustering (SuperSCC), a novel algorithm that utilizes machine-learning models to discover cell identities and gene modules from multiple datasets without the need of data integration. Of note, SuperSCC can be implemented both in cell lineage and cell state level, thereby building the hierarchy of cell programs with specific cell identity and gene modules. Such information has the great potential to identify the shared rare populations across datasets regardless of batch effect and benefits label transfer for mapping cell labels from reference to query. We used SuperSCC to perform atlas level data analysis on more than 90 datasets and build a cell state map of complex tissue in healthy and diseased stages, such as human lung. We show that SuperSCC outperforms existing approaches in identifying cellular context, has better annotation accuracy, and outlines gene modules that indicate conserved immune cell status in lung microenvironments.

## Introduction

Single-cell RNA sequencing (scRNA-seq) has revolutionized biological research by providing insights at the individual cell level, offering a detailed view of transcriptome regulation^1–3^. As the volume of publicly available single-cell datasets rapidly expands ^4,5^, there is an urgent need to integrate these datasets to construct comprehensive cellular atlases, which are essential for a deeper understanding of complex biological systems. Such integrative approaches allow researchers to explore cellular hierarchies and gene expression dynamics across a range of conditions, including health and disease.

Despite the advancements in scRNA-seq technologies, current methods for data integration face significant challenges. A critical issue is how to integrate datasets from different sources while preserving biologically relevant variability and avoiding overcorrection of key signals^6^. The necessity for batch correction prior to downstream analyses, such as clustering and cell type annotation, can introduce biases and obscure biologically meaningful variations, complicating the interpretation of integrated datasets^7^.

In addition, standard approaches for clustering and marker gene detection often fail to capture the inherent hierarchical structure of cell types and states. For instance, broad cellular categories like lymphoid cells and more specific subpopulations such as CD8^+^ T cells exist at different levels of cellular hierarchy. Yet current tools, including Seurat^8^, Scanpy^9^, and SCMarker^10^, struggle to accurately distinguish these levels. These hierarchical relationships are critical for understanding functional diversity within tissues. While recent methods like Spectra^11^ have addressed some challenges by identifying gene programs without batch correction on large atlases, they lack the ability to construct hierarchical structures and require prior knowledge in the form of gene sets.

To address these limitations, we developed SuperSCC (Super Single Cell Clustering), a novel computational framework that enhances scRNA-seq analysis by enabling hierarchical clustering, marker gene detection, and robust label transfer across multiple datasets without the need for batch correction. SuperSCC performs iterative clustering, gene module detection, and cluster merging to construct cell type hierarchies and gene modules at multiple levels. We applied SuperSCC to large-scale datasets, including human lung tissues and immune cell atlases, across diverse conditions and diseases. Our results demonstrate superior performance in identifying cellular contexts, annotating cell states, and discovering conserved gene modules compared to current state-of-the-art methods. Notably, SuperSCC uncovered several rare and conserved immune cell states, some of which are linked to immunotherapy efficacy, underscoring its potential to reveal novel insights into disease biology.

## Methods

The SuperSCC algorithm consists of three major aspects. First, a supervised feature selection process leveraging filter-, embedded- and wrapper-based methods is implemented to capture the most informative features in the dataset and this is the cornerstone of SuperSCC and refers to Detector here. Second, a cluster-aware clustering is built on the combination of graph-based clustering with Detector, resulting in highly similar clusters merging in different levels and hence establishment of gene module and cell identity hierarchy. Third, training and modeling reference dataset with known cell labels and key features found by Detector and then assignment of cell labels in the reference dataset to query dataset via calculating the similarity of feature loadings between them. The details of each of those steps have been outlined below.

### Detector - a supervised feature selection process

Detector starts with a cell-by-gene matrix with an additional column containing the cell labels from the data generator or cluster labels from any clustering algorithm. The initial feature selection is done by removing low-variance features in the dataset and the remaining features are called seed features. Such variance cut-off is the average or median value of variances across all features in the dataset. Then, in the case of filter-based category, seed features process with correlation filter through conducting one of F test, Chi-square test or mutual information (MI) analysis, thereby ranking features based on corresponding statistics or discarding away features being independent of class and irrelevant for classification.

The F statistics is denoted as:

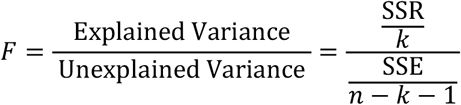

Where SSR is the sum of squares due to regression (explained variance). SSE is the sum of squares due to error (unexplained variance). k is the number of predictors (independent variables). n is the number of observations.

The *χ*^2^ statistics is denoted as:

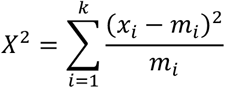

Where X_i_ is the observed frequency and m_i_ is the expected frequency for category i respectively. The MI statistics is denoted as:

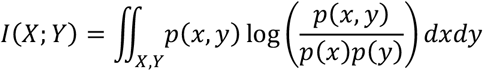

Where p(x, y) is the joint probability density function of X and Y, p(x) and p(y) are the marginal probability density functions of X and Y.

With intersection mode, for F test or Chi-square (*χ*^2^) test, only seed features with statistical significance (*p* value < 0.05) are kept while seed features with non-zero evaluation score remain as doing mutual information. With ensemble mode, the seed features are ranked by the specific statistic in a descending order.

Then, in the case of embedded-based categories, the seed features are utilized to train an induction model such as support vector machine (SVM), logistic regression (LR) or random forest (RF) and then are scored based on importance weights obtained from model fitting. As the model is SVM or LR, seed features are scored by coefficient attributes, which are the best model parameters for classification.

When SVM, the loss function of SVM model is defined as:

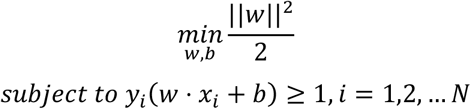

Where ∥w∥ represents the magnitude of the model parameters w. As the whole formula reaches the minimum, w indicates the best model parameters and is aggregated to score each feature. With ensemble mode, seed features are ranked by aggregated absolute *w* in a descending order. With intersection mode, only seed features with non-zero *w* remain after L1 regulation.

When LR, the loss function of LR model is defined as:

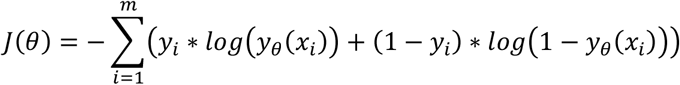

where θ is a set of parameters, m is the number of observations, y_i_ is the real label of observation and y_θ_(x_i_) means the estimation of observation i is based on θ model parameters. As J(θ) reaches the minimum, θ indicates the best model parameters and is aggregated together to score each feature. With ensemble mode, seed features are ranked by aggregated absolute θ in a descending order. With intersection mode, only seed features with non-zero θ remain after L1 regulation.

As model is RF, seed features are scored by feature importance (also known as mean decrease in impurity) attribute which are defined as:

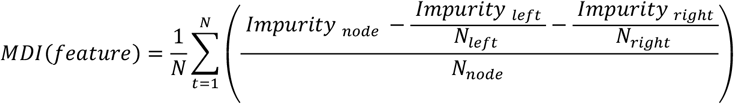

where N is the total number of trees in the forest, Impurity_node_ is the impurity of the parent node before the split, Impurity_left_ and Impurity_right_ are the impurities of the child nodes after the split using the feature, *N*_left_ and *N*_right_ are the number of samples in the left and right child nodes respectively and *N*_node_ is the total number of samples in the parent node. With ensemble mode, seed features are ranked by feature importance in a descending order. With intersection mode, only seed features remain as their feature importances above 1/N_sf_ where N_sf_ is the total number of seed features.

Furthermore, in the case of wrapper-based category, recursive feature elimination (RFE) algorithm combines one of induction models mentioned above to rank features based on importance weights or remove the most insignificant features iteratively until the number of features reaches the expected size. Given the risk of overfitting, the RFE algorithm runs on the different cross-validation splits, thereby automatically tuning the expected size of selected features. The number of selected features is equal to the number of features maximizing the cross-validation score. With ensemble mode, seed features are ranked by importance weights calculated by induction model in a descending order. With intersection mode, only surviving features after accomplishing the RFE algorithm remain.

Finally, with integrating results from all three categories, the selected feature set is the intersection of feature subsets from each feature selection category while the comprehensive feature ranking is the geometric mean of feature rankings from each feature selection category.

### Cluster-aware clustering process

This process starts with cell-by-gene matrix, followed by applying Leiden algorithm on k nearest neighbors (KNN) graph to obtain the raw cluster (Rcluster) labels. Then, the highly variable genes (HVGs) that best explain variation in the dataset and also highly correlate with Rcluster are found by Detector. To capture the representatives of each Rcluster from HVGs pool, Detector was implemented in one-vs-rest mode (referring to SuperOVR), in which all other Rclusters excluding the interest Rcluster are treated as a big group. Two similarity matrices are generated by calculating the pairwise jaccard index or distance correlation.

The Jaccard index is defined as:

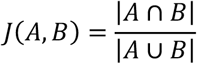

where A and B mean the key gene sets (found by SuperOVR) from two clusters between comparisons.

The distance correlation^12^ is denoted as:

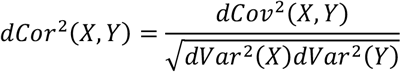

where X and Y represent the normalized expression of shared representatives from two clusters, dCov and dVar means the distance covariance and distance variance respectively.

The product of those two similarity matrices is the score determining whether two compared clusters should be merged. Only two clusters between comparison with above 0.1 score would be merged. As there are multiple satisfied cluster pairs with ties, those two compared clusters with the highest score would be merged. After that, the merged cluster labels are treated as the cell label input for Detector and run this process recursively until no merging candidates exist. The final merged clusters are theoretically cell lineage clusters and refer to M clusters here. Then, in each M cluster, we repeat above procedure with three minor changes including 1) changing the default resolution from 1 to 0.2 when applying Leiden algorithm, 2) the best representatives of each sub clusters not from HVGs pool but from all features in the dataset and 3) merging clusters with less than 10 positive markers. In this manner, cell state clusters, referring to F clusters are obtained and hence enable building the hierarchy of cell identity and gene module.

### Label transfer process

This process starts with a cell-by-gene reference matrix containing known cell labels, which processes with Detector to find the HVGs. Then, the reference matrix would be filtered to only keep HVGs. Such data subset executes grid search on cross-validation splits to evaluate the best specified model parameters regarding different induction models. In default, parameters (“C”, “kernel”) for SVM model, parameters (“penalty”, “C” and “multi_class”) for LR model, and parameters (“n_estimatorss”, “criterion” and “max_features”) for RF model would be searched. Otherwise, customized parameters for the corresponding model found on scikit-learn official website can be used for parameters optimization. Then, the induction model uses the best model parameters for model training on a reference dataset. As for prediction on query dataset without cell labels, the query dataset also filters to retain the HVGs found in the reference dataset. If the HVGs are not in the query dataset, those genes would be padded with zero to ensure the feature dimension being matched between query and reference dataset. Then, mapping the features of each data point in the query dataset to feature space used for model training gets the predicted labels. In brief, as the model is SVM, the decision function assigns novel data points to labels in the nearest support vector. As the model is LR, the decision function assigns novel data points to labels of the highest probability after sigmoid transformation. As the model is RF, the decision function assigns novel data points to labels being voted most by all independent decision trees.

### Gene module detection

As described previously^13^, the defining of gene modules follows a Jaccard similarity-based approach. In brief, for each M cluster or F cluster across all collected datasets, we focus on those genes with log2 fold change above 1 and being exclusively expressed in the interest M/F cluster and then rank them by importance weights obtained from SuperOVR and consequently get a list of top 50 genes for gene module detection. Each gene set is compared to all other gene sets to evaluate the degree of gene overlap between gene sets. Given overlapping instances of at least 10 genes, the compared two gene sets with the maximal overlapping rate is treated as the founder of a new gene module. As the number of overlapping gene sets with at least 10 genes exceeds 5 cases, the gene set with the highest overlapping rate to the founder is added. To ensure each gene module always consists of 50 members, other genes besides shared ones are ranked by importance weights and genes with top importance weights are kept. Notably, for shared genes between comparisons, higher importance weight remains for next round gene module updating. Such a process is then repeated until no satisfied candidate that can be added to the gene module exists. After accomplishing the first gene module, the remaining gene sets process with above procedure again by searching for the founder of gene module, updating the founder via adding the gene set with maximal overlap to the founder as well as completing the gene module list to 50 genes according to importance weights in the pool of gene sets, thereby forming the second gene module and so far. Notably, in this manuscript, we only used gene modules consisting of at least three gene sets from different datasets for downstream analysis and finally got 15 MGMs and 65 FGMs.

### Data curation

The detail of the dataset used in this manuscript was shown in Supplementary Table 1. For dataset selection, we first generated a list of potentially relevant studies through search on PubMed, Gene Expression Omnibus (GEO), CELL x GENE and EMBL-EBI. Each dataset was examined whether raw sequencing files/count matrix could be publicly accessed. If not, it would be filtered. Besides, those with atlas-level data and expert annotation were prioritized.

Finally, we collected 99 datasets for downstream analysis. For data preprocessing, if the study provided a quality-control (QC) counts matrix, we directly used that, otherwise, we followed the filtering rule in the source paper of the corresponding dataset to remove low-quality cells and low-abundance features. After QC, per dataset we obtained 20,000 cells subset with same cell type frequency (Python package: sklearn^14^[v: 1.2.2], train_test_split) compared to the original one and then did log normalization (feature counts for each cell are divided by the total counts for that cell and multiplied by the 10000, followed by natural-log transformation (R package: Seurat [v: 4.3.0], NormalizeData).

### Running SuperSCC

Log-normalized counts matrix was treated as the input for SuperSCC and ran in default settings to obtain the M (SuperSCC, global_consensus_cluster) or F clusters (SuperSCC, sub_consensus_cluster) and the corresponding representative cluster markers in each dataset. Such M/F clusters and relevant representative markers (top 50) were used for downstream clustering benchmarking and building the conserved gene modules hierarchy based on the aforementioned gene module detection method. Log-normalized counts matrix plus cell type labels (provided by data generator) fed to SuperSCC find_markers_ovr function to get top 20 markers per cell type per dataset that were finally used for feature selection benchmarking. As for executing SuperSCC label transfer pipeline, data fed to SuperSCC feature_selection function to capture the overall best representative features followed by model training and label prediction using SuperSCC model_training and predict_label function respectively. Such label transfer pipeline was run with different input data: 1) counts matrix with cell type labels was split into train and test subset, accounting for 70% and 30% original data respectively. The train subset was for the training model while label prediction was finished on the test subset. This refers to the ‘self-validation’ category. 2) model training and label prediction were done on separate dataset. This refers to the ‘cross-validation’ category. The SuperSCC predictions are retained for label transfer benchmarking.

Eighteen collected datasets were used for above-mentioned benchmarking in feature selection and clustering (Supplementary Table 1). Nine and six collected datasets for label transfer in self-validation and cross-validation respectively. The reference dataset used for label transfer model training in cross-validation was produced by randomly selecting cells multiple datasets (Supplementary Table 1)

### Running Scanpy and Seurat

Both Scanpy and Seurat were performed by default processing steps (normalization, finding highly variable genes, principle canonical analysis (PCA), clustering and finding markers) to obtain clusters and their markers. The key differences of clustering between Scanpy and Seurat are two points: 1) Seurat requires scaling data before PCA but not for Scanpy; 2) Seurat applies Louvain algorithm with default 0.8 resolution for community detection within neighborhood graph while Scanpy uses Leiden algorithm with default 1 resolution for that. Clusters from both tools were used for clustering benchmarking. Similarly, Scanpy (Python package: Scanpy [v: 1.9.8], rank_genes_groups) and Seurat (R package: Seurat [v: 4.3.0], FindAllMarkers) were also applied on data grouped by cell type to retain top 20 markers per cell type per dataset, followed by feature selection benchmarking.

### Running SC3, CIDR, DUBStepR, SingleR and SingleCellNet

SC3^15^ (v: 1.30.0), CIDR^16^ (v: 0.1.5) and DUBStepR^17^ (v: 1.2.0), SingleR^18^ (v: 2.4.1) and SingleCellNet^19^ (v: 0.1.0) were run in default settings according to their user guide. Of note, since SC3 requires users to determine the k to infer clusters, we set the k to the number of M clusters detected by SuperSCC when running SC3. The clusters from SC3, CIDR and DUBStepR were used for clustering benchmarking. As running SingleR and SingleCellNet, both tools were run in self-validation and cross-validation modes, same as what was done on the SuperSCC label transfer pipeline. The predictions from SingleR and SingleCellNet were used for label transfer benchmarking.

### Feature selection, clustering and label transfer benchmarking

For feature selection benchmarking, we utilized large language models^20^ (LLMs) to assess which gene sets provided by different algorithms (SuperSCC, Scanpy and Seurat) offer the best representation of the corresponding cell type. The ideal prompt for gene sets evaluation was produced by prompt engineering and iterative refinement (Extended Data Fig. 1). The evaluation output includes two scoring metrics: relevant gene ratio and biological relevance score. The former indicates the proportion of genes associated with signatures of relevant cell type to the total number of input genes while the latter derives a relevance score associated with cell type specific signatures based on Gene Ontology (GO) and KEGG collections for each gene set. Finally, we averaged relevant gene ratio/biological relevance scores across cell types to obtain the overall score reflecting the performance of capturing biologically meaningful features per dataset per algorithm. Besides, we also show the trend between biological relevance score and cell type frequency to highlight the efficacy of each algorithm under low (< 5%) and extremely low cell type (< 15%) frequency.

For clustering benchmarking, clusters detected by five different algorithms (SuperSCC, Scanpy, Seurat, CIDR, DUBStepR) and ground truth cell type labels were used for calculating four scoring matrices including adjusted mutual information (AMI), adjusted rand index (ARI), normalized mutual information (NMI) and Fowlkes-Mallows index (FMI). For label transfer benchmarking, predicted labels from three tools (SuperSCC, SingleR, and SingleCellNet) were utilized, either through self-validation or cross-validation, along with ground truth cell type labels, to compute eight scoring matrices such as accuracy score (ACC), homogeneity score (Homog), recall score, precision score (Prec), completeness score (Compl), Matthews correlation coefficient (MCC), F1 score and V measure score (VM).

### Gene module enrichment/conservation analysis

We mainly used signatures from Gene Ontology collections (C5.GOBP, C5.GOCC and C5.GOMF). Hypergeometric test (R package: clusterProfiler^21^ [v: 4.10.1], enrichGO) was applied to test whether provided gene modules enrich on corresponding signatures. Signatures with false discovery rate (FDR) < 0.05 were considered as significant. 15 MGMs and 65 FGMs were further grouped by functional similarities into 6 and 11 hallmarks respectively. In addition, overlapping rates for comparisons among 238 SuperSCC M clusters’ gene sets or 631 SuperSCC F clusters’ gene sets based on their top 50 genes within 99 datasets across 20 tissue types were calculated. Furthermore, we counted the number of gene sets contributing to GMs across datasets and then sum those numbers based on hall markers to denote the frequency of each hall marker in a specific tissue type or across tissue types.

Then, in each module, we obtained the representative cell type via digging its contributor (the clusters with gene set devoting to the module) to that module. Then, we calculated gene module activity score (R package: Seurat [v: 4.3.0], AddModuleScore) and counted the frequency of conservation events in which dots with high activity score are consistent with expected cell type across 99 test datasets. Overall conservation frequency and conservation frequency per tissue were used to show module conservation at different levels.

### Connection analysis between MGMs and FGMs

To assess the connection between MGMs and FGMs, we combined hierarchy clustering and GMs information together to bridge modules in different levels and functionalities. We first obtained contributor F clusters for all 65 FGMs and traced back their M clusters as well as MGMs in the following approach: if F cluster A contributing to FGM1 was child cluster of M cluster B, we examined whether M cluster B contributed to any MGMs and if yes recorded the MGM name, otherwise unknown source remained. Then, for each FGM, we calculated the frequency towards different MGMs or unknown source. Finally, such pairwise MGMs to FGMs chart with frequency number, contributor M/F clusters and also functional hallmarks of corresponding modules were treated as input for Sankey plot visualization (Python package: Plotly^22^ [v: 5.22.0], Sankey).

### Hierarchy clustering, Pearson correlation and differential gene expression analysis

For hierarchy clustering, Jaccard similarity among 65 FGMs was calculated, followed by pairwise distance computation (R package: base [v: 4.3.1], dist(method = “euclidean”)) and final clustering step (R package: base, hclust(method = “average”)).

For Pearson correlation, in dataset Dong_2020_1 we calculated FGM32 and MGM4 activity scores and extracted both scores in cells of F cluster 14 for correlation analysis (R package: base, cor(method = “pearson”)). Then, a hypothesis test was performed to see whether there was a positive correlation between scores (R package: base, cot.test(method = “pearson”)). The *p* value < 0.05 mean scores A positively correlated with scores B. Similarly, in the dataset Laughney_2020, we did correlation analysis between F cluster 6 and other F clusters based on the frequency of patients.

For differential gene expression analysis, a t-test was done to compare the gene expression between F cluster 14 and F cluster 18 in dataset Dong_2020_1 or between stress B-associated T cells and other T cells in dataset Laughney_2020. Genes with FDR-corrected *p* value < 0.05 and |log2foldchange| > 1 was considered a differential expression.

### Visualization

All graphs in this manuscript were drawn by using R package ggplot2^23^ (v: 3.5.1) except the specified statement.

## Results

### The workflow of SuperSCC

SuperSCC is a generalized framework for multiple tasks on single-cell transcriptomic data including informative features extraction, cluster-aware clustering and label transfer. Firstly, the framework uses a combination of machine learning approaches to identify informative features with the best representation of the data and significant correlation with cell types/clusters (Fig. 1a, top; Extended Data Fig. 2a). Such supervised methods are able to discover abundant small differences in gene expression between different cell types, benefiting cell clustering and label transfer. Given that, SuperSCC enables cluster-aware clustering that avoids over clustering or under clustering and also allows the construction of cell identity hierarchy via merging similar clusters in different levels (Fig. 1a, middle left; Extended Data Fig. 2b). Furthermore, SuperSCC could map cell labels from reference to query (Fig. 1a, middle right; Extended Data Fig. 2c). To run SuperSCC, it requires a normalized cell-by-gene count matrix with cell/cluster labels as input and outputs a set of profiles for different tasks. For informative features extraction, SuperSCC provides a small pool of genes maximally representative for a given dataset that would be utilized for further cluster-aware clustering and label transfer. For cluster-aware clustering, SuperSCC gives the broad clustering (cell lineage level) and finest clustering (cell state level) and also the informative genes for corresponding identified clusters. For label transfer, the predicted class and the class probabilities for query would be returned.

**Fig 1.**
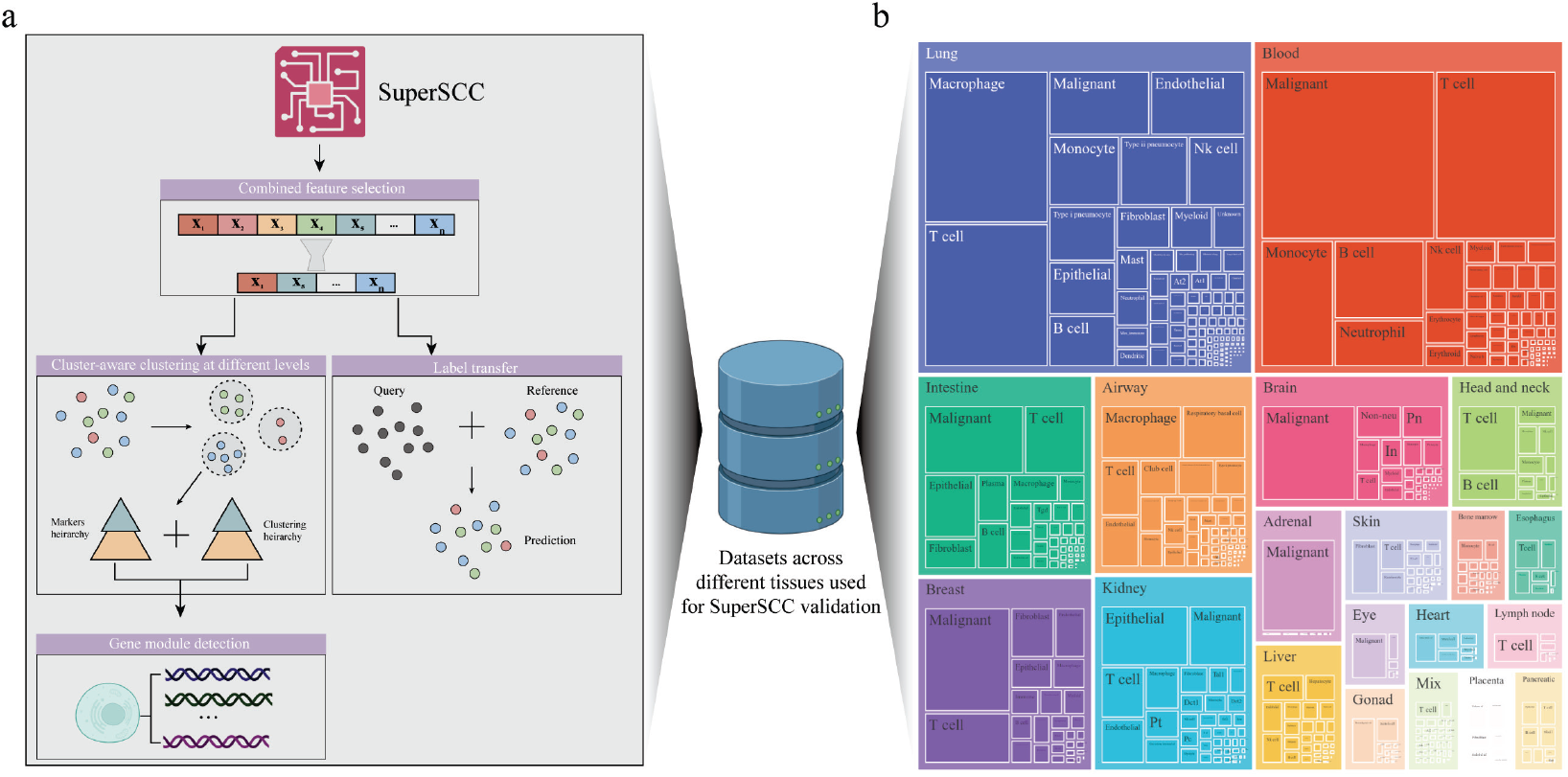
SuperSCC covers multiple analysis tasks on scRNAseq. **a**, The schematic overview of the SuperSCC framework. SuperSCC combines machine-learning feature selection approaches to capture the informative features within single-cell transcriptome data, facilitating cluster-aware clustering and label transfer. Cluster-aware clustering provides clustering and corresponding markers at different levels, thereby enabling building cell program hierarchy. Such hierarchical information could be further utilized for detection of conserved gene modules across datasets without the need of data integration, making ‘analysis-then-merge’ strategy. Label transfer here is reference-based automatic cell annotation, whereby the cell label in the reference can be assigned to unlabeled cell in the query with similar gene expression profile. **b**, Treemap plot showing the dataset composition across tissues and cell types. The outer rectangles reflect the number of datasets per tissue, the inner rectangles correspond to the frequency of each cell type.

Two key features make SuperSCC have greater performance regarding above-mentioned scRNAseq tasks than existing tools. First, with the advantage of cluster-aware clustering, SuperSCC overcomes the clustering bias when implementing graph-based clustering method under an inappropriate resolution parameter^24^, thereby mitigating over/under clustering and further recovering cell ontology tree, providing biologically meaningful cell program in hierarchy. Second, such hierarchical cell program information can be further used for defining conserved gene modules (Fig. 1a, bottom) and hence shared rare populations without the need of data integration or batch effect correction across datasets, which may compromise latent biological differences. To help interpret the result and compare gene modules, we integrate LLM agent into SuperSCC to facilitate biological knowledge discovery, which can also be used as a benchmarking tool. Finally, we used SuperSCC to do atlas level data analysis on almost 100 datasets among different tissue types and disease states (Fig. 1b; Supplementary Table 1).

### SuperSCC has great capability for multiple scRNAseq analysis tasks

We manually examined the quality of cell annotation and selected 18 datasets with expert-annotated cell labels out of the dataset pool used in this manuscript. We applied SuperSCC and other widely used tools on those datasets,then compared their performance on various tasks. First, we employed LLM to assess the performance of SuperSCC, Scanpy^9^ and Seurat^8^ on capturing informative features (Fig. 2a; see Methods) via two scoring metrics including relevant gene ratio and biological relevance score. The former denotes the proportion of genes associated with signatures of relevant cell type, while the latter indicates a relevance score associated with cell type specific signatures. SuperSCC (SuperSCC: 0.486 ± 0.0169) and Scanpy had comparable relevant gene ratios, which is higher than Seurat (Fig. 2b, left; Extended Data Fig. 3a). Similar observation was also found in biological relevance scoring. SuperSCC got the highest score (0.710 ± 0.0151) while Scanpy and Seurat came in second and third respectively (Fig. 2b, middle). Moreover, when cell type frequency ranged from low (∼5%) to medium (∼15%), SuperSCC (0.791 ± 0.0186) showed much greater performance of capturing informative features (Fig. 2b, right; Extended Data Fig. 3a) than Scanpy and Seurat.

**Fig 2.**
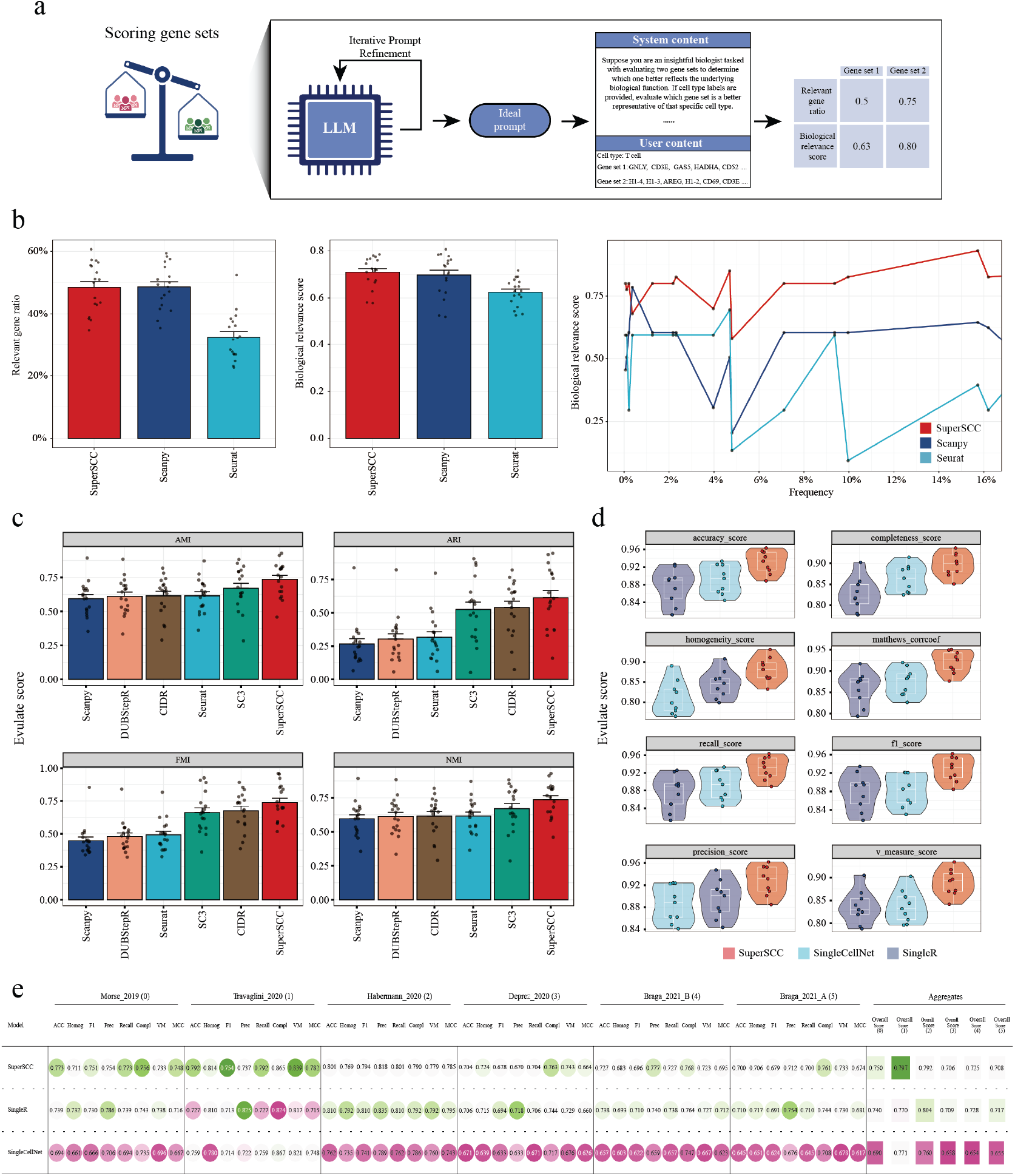
Evaluation of SuperSCC performance on multiple scRNAseq analysis tasks. **a**, The schematic overview of employing a large language (LLM) model assessing which input gene set is a better representative of the provided cell type. The prompt was produced by prompt engineering and iterative refinement (see Methods). **b**, Relevant gene ratio (left), biological relevance score (middle) and biological relevance score trend along with cell type frequency (right) of three comparing algorithms SuperSCC, Scanpy and Seurat on eighteen testing datasets. Bar plots were presented as mean ± SEM, with each dot representing the aggregated overall score on the individual testing dataset. **c**, AMI, ARI, FMI and NMI scores of SuperSCC, Scanpy, Seurat, CIDR and DUBStepR on eighteen testing datasets. Bar plots were presented as mean ± SEM, with each dot representing the aggregated overall score on the individual testing dataset. **d**, ACC, Homog, Recall, Prec, Compl, MCC, F1 and VM scores of SuperSCC, SingleR and SingleCellNet on nine testing datasets under self-validation whereby model training and prediction were done on the same dataset. Boxes and lines indicated interquartile range (IQR) and median; whiskers were 1.5x IQR, with each dot representing the overall score on the individual testing dataset. Violins displayed the data distribution. **e**, same with d but in cross-validation in which model training and prediction were done on separate dataset. Color range displayed the performance per scoring matrix per algorithm under the corresponding dataset. The greener the dot, the higher the score. Aggregated scores were the weighted average of eight scores per algorithm per dataset.

Furthermore, we also evaluated whether SuperSCC outperforms other tools on recovering broad cell ontology. To do this, we measured similarity between clusterings of each method and ground-truth cell labels provided by the data generator. SuperSCC showed the best performance on more than 60% (11 out of 18) validation datasets (Extended Data Fig. 3b) and its overall score ranked top 1 in all four evaluation metrics AMI (0.736 ± 0.0297), ARI (0.613 ± 0.0540), FMI (0.738 ± 0.0329) and NMI (0.736 ± 0.0297) (Fig. 2c). Notably, SuperSCC got 2 times higher overall ARI and 1.5 times higher overall FMI score than methods Seurat and Scanpy. Besides, SC3^15^ clustering in which k was determined by the number of M clusters searched by SuperSCC (see Methods) ranked in second in AMI and NMI scoring and its ARI and FMI scores were only slightly lower than CIDR^16^ but still higher than both Seurat and Scanpy (Fig. 2c).

Moreover, we further benchmark the performance of SuperSCC on label transfer application within self-validation and cross-validation categories which are classified by whether training and testing were performed on the same data (see methods). With self-validation, all tested algorithms performed great and had above average 0.8 overall scores in total eight scoring metrics (Fig. 2d). Of note, SuperSCC always showed the best performance with acquiring the highest scores in ACC (0.933 ± 0.0608), Homog (0.881 ± 0.0566), Recall (0.933 ± 0.0608), Prec (0.931 ± 0.0654), Compl (0.899 ± 0.0597), MCC (0.926 ± 0.0584), F1 (0.933 ± 0.0608) and VM (0.893 ± 0.0692) (Fig. 2d; ; Extended Data Fig. 3c). SingleCellNet^19^ and SingleR^18^ had comparable performance. SingleCellNet showed higher scores in ACC, Compl, MCC and recall and VM than SingleR while an opposite trend was observed in F1, Homog, and Prec (Fig. 2d). With cross-validation, both SuperSCC and SingleR had comparable performance on six tested datasets apart from the second in which SuperSCC got ∼ 0.3 overall score higher than SingleR (Fig. 2e). Detailed evaluation scores per algorithm per scoring matrix could be found in Supplementary Table 2.

### SuperSCC discovers conserved and hierarchically organized gene modules across tissue atlases

Traditional single-cell analysis pipelines rely on data integration prior to downstream clustering, risking the loss of nuanced biological signals during batch correction ^7,8^. SuperSCC circumvents this limitation through an “analysis-then-merge” framework, first resolving cell identity hierarchies within individual datasets to detect conserved gene modules (GMs) and subsequently aggregating these findings across tissues (Fig. 3a). By leveraging hierarchical clustering, SuperSCC distinguishes broad cell identity-associated modules (M clusters’ GMs; MGMs) from fine-grained state-specific modules (F clusters’ GMs; FGMs) on 99 datasets spanning 20 tissues. At its broadest resolution, SuperSCC defines 15 major gene modules (MGMs) (Supplementary Table 3) representing core cellular programs (e.g., immune response, angiogenesis), organized into six functional families (Fig. 3b). These MGMs are enriched in lineage-defining pathways and exhibit tissue-specific dominance, with immune-response MGMs prevailing in diverse tissues (54.6% of all MGMs), while cytoskeleton-associated MGMs uniquely characterize respiratory airway datasets (12%, Fig. 3b, right). At finer resolution, 65 sub-modules (FGMs) (Supplementary Table 3) delineate cell-state-specific programs, including novel processes absent in MGMs, such as antioxidant/stress responses (10% of FGMs) and energy synthesis (Fig. 3c; Extended Data Fig. 5a). Critically, FGMs often refine MGMs: cell-cycle FGMs, prevalent in brain datasets (52%), elaborate on cell-development MGMs, while angiogenesis MGMs (24.8%) expand into immune-linked sub-modules under FGM level (Fig. 3c, Extended Data Fig. 5a).

**Fig 3.**
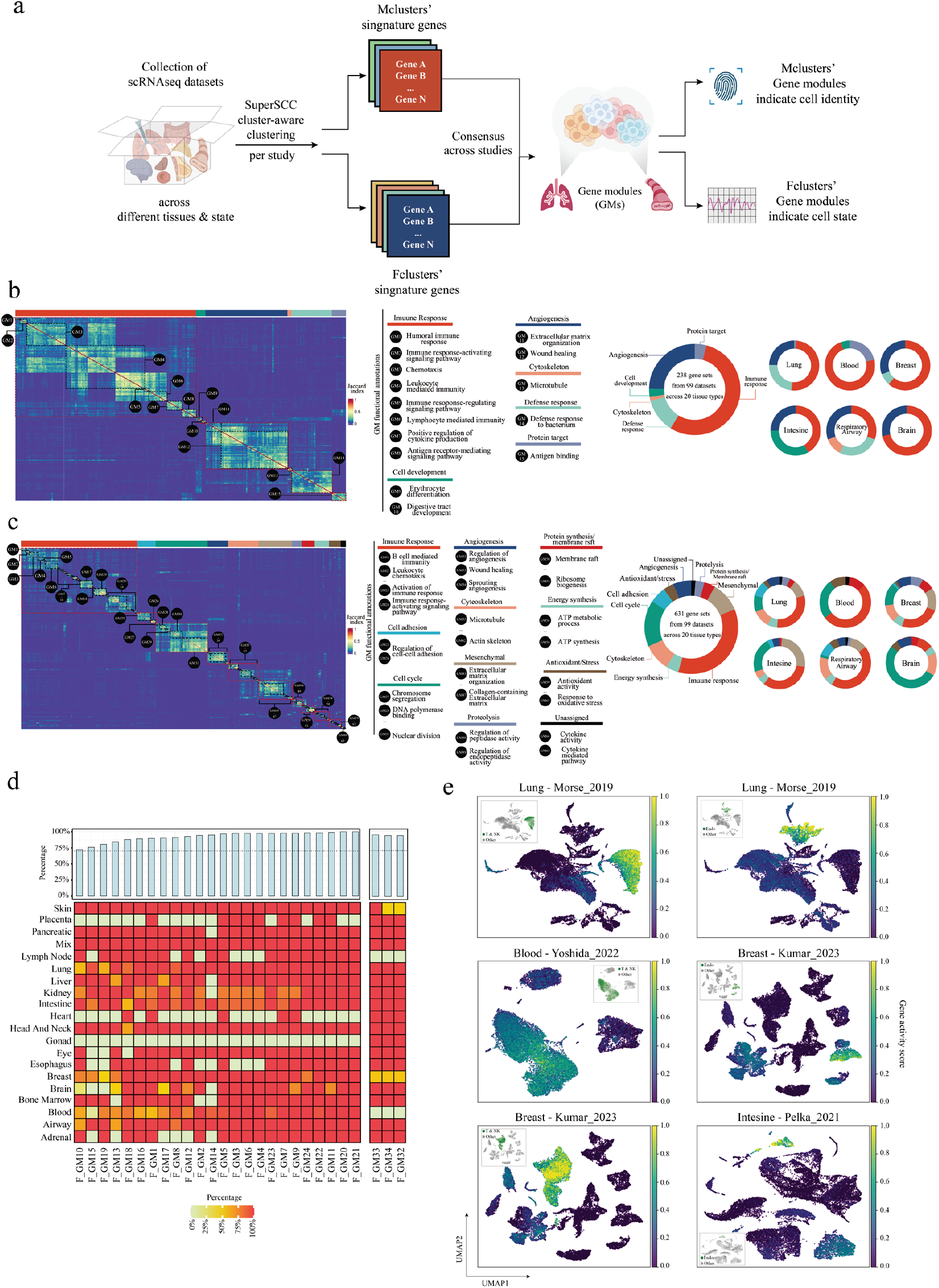
SuperSCC discovers conserved gene modules across datasets and tissue types without the need for data integration. **a**, The schematic overview showing SuperSCC employee hierarchy cell programs from cluster-aware clustering on each individual dataset to find consensus gene modules (GMs) across datasets and tissue types. **b**, Heatmap showing Jaccard similarity scores for comparisons among 238 SuperSCC M clusters’ gene sets based on their top 50 genes within 99 datasets across 20 tissue types. Gene sets were ordered by clustering and grouped into GMs (circled by black dashed lines) and families of GMs with similar functionalities (marked by red dashed lines and top column annotations). GMs were numbered and labelled. Lists of pathways enriched on GMs were shown by side. The bigger circle on right reflected the frequency of six GM families across all datasets and tissue types, while smaller circles focused on the GM families’ frequencies in a particular tissue type. **c**, Same as b but under F cluster level. Eleven GM families’ frequencies were shown. **d**, Conservation frequency of each immune response (left) or angiogenesis (right) F cluster’s GM (FGM) in (columns) each tissue type (rows). The top annotation bar presented the aggregated average conservation frequency per GM across all tissue types. Color range associated with frequency. The redder the box, the higher the frequency. **e**, the outer UMAP showing the activity score of immune response FGM20 and angiogenesis FGM33 in d in different tissue types in different studies. The more yellow the dot, the higher the score. The inner UMAP shows the region of expected cell type (green dots) that were supposed to express the highest activity score of the corresponding FGMs. The cell label was provided by the data generator.

Strikingly, 24 immune-response and three angiogenesis FGMs were conserved in >70% of datasets, with activity localized to expected cell types (e.g., immune FGMs in T cells, Fig. 3d–e), underscoring SuperSCC’s capacity to resolve context-independent signatures. Such gene module conservancy was also observed in MGMs (Extended Data Fig. 4). This hierarchical modularity—broad MGMs anchoring core functions and FGMs capturing dynamic states— provides a scaffold to interrogate functional cross-talk between cellular programs.

### Hierarchical gene modules reveal cross-compartment immune regulation

The hierarchical relationship between MGMs and FGMs enables the discovery of dynamic gene functionality across biological scales. Unlike static module-detection methods^11,13,25^, SuperSCC reveals how broad cellular programs bifurcate into specialized states, particularly within immune contexts. We demonstrate this by exploring two phenomena: stress adaptation in immune cells and immune-activation signatures in non-immune populations.

Within immune-response MGMs, non-immune FGMs emerged, such as antioxidant/stress (FGM54/FGM55) and energy synthesis (FGM62/FGM63) modules, enriched in heat shock genes (Fig. 4a–c). These FGMs distinguished stressed immune subsets (e.g., activated B/T cells, macrophages) through upregulated heat shock protein expression (Fig. 4c; Extended Data Fig. 5b), illustrating how SuperSCC disentangles functional diversification within immune lineages. Such observations were consistent with previous study in which stress T cells were characterized by *HSPA1A*/*HSPA1B*^*26*^. However, SuperSCC’s findings extended the stress response state beyond T cells to other immune cells and also highlighted novel relevant heat shock genes such as *HSPB1, HSPD1, HSPH1, HSPA4* and *HSPA6*, which were reported to be involved in immune response^27–29^ but less being treated as indicators of stress immune cells. Furthermore, we examined the frequency of stress B cells (F cluster 6) under different clinical conditions in Laughney_2020 and found stress B cells remarkably decreased in treated lung cancer patients compared to the naive patients (Fig. 4d). Interestingly, correlation analysis based on the frequency of cells in each patient showed a strong association between F cluster 6 and F clusters 2, 4 and 30 which were T cells and referenced to stress B cells-associated T cells here (Fig. 4d). Further DE analysis comparing stress B cells-associated T cells with other T cells found the up-regulated heat shock genes, suggesting that those stress B cells-associated T cells were also in stress response state (Fig. 4e).

**Fig 4.**
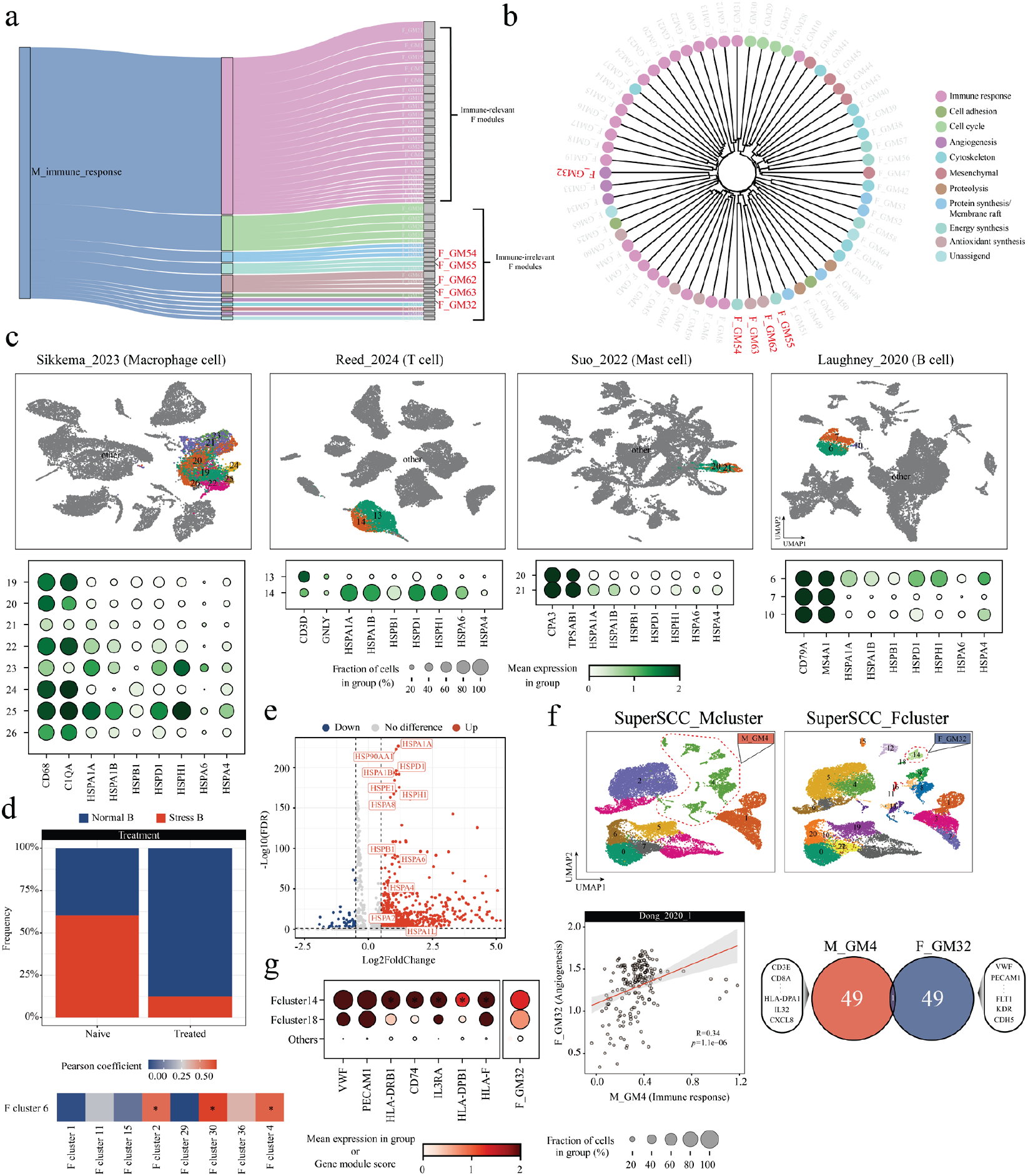
SuperSCC-derived gene modules find rare cell states in multiple immune cell types and immune-active non-immune cells. **a**, Sankey plot showing the connection between immune response MGMs and FGMs in different functions. Numbered FGMs were shown. Immune-relevant GMs were in pink while immune-irrelevant GMs were in other colors which represented the functions shown in color legend in b. Interesting FGMs discussed in the text were highlighted in red. **b**, Hierarchy clustering of 65 FGMs based on their pairwise Jaccard similarity. Different colors revealed different functionalities of each FGMs. Interesting FGMs discussed in the text were highlighted in red. **c**, UMAP (top) showing the SuperSCC F clustering with only highlighting the clusters of the focused immune cell types in datasets Lauughney_2020 (B cell), Reed_2024 (T cell), Suo_2022 (Mast cell) and Sikkema_2023 (Macrophage cell). Dot plots (bottom) showing the classical markers of focused immune cell types in each dataset and heat shock genes. The size of dots reflected the fraction of cells in corresponding clusters. The color range represented the average expression of genes in that cluster, whereby the greener the color, the higher the expression. **d**, Bar plot showing the frequency of stress B cell in naive and treated patients on Laughney_2020. Color red and blue represented stress and normal B cell respectively. Heatmap showing the Pearson correlation on Laughney_2020 regarding the frequency of patient samples between F cluster 6 and other F clusters containing cells from all patients. The redder the box, the higher the frequency. The asterisk indicated significant correlation with *p* value < 0.05 from correlation test. **e**, Volcano plot showing the differential expression (DE) genes by comparing stress B cell-associated T cells versus normal T cells. The DE genes was determined by |log2FoldChange| > 0.5 & - log10(FDR) < 0.05. Upregulated heat shock genes in stress B cell-associated T cells were highlighted. **f**, UMAP showing the SuperSCC M (top left) and F clustering (top right) on Dong_2020_1. The dashed lines circled the M cluster 14 and F cluster 18, which contributed to MGM4 and FGM32 respectively. Scatter plot (bottom left) showing the correlation between MGM4 and FGM32 activity scores in cells within F cluster 14 in Dong_2020_1. Each dot represents one cell. Linear regression line with 95% confidence interval was shown. Pearson coefficient and *p* value were provided by Pearson correlation test. Venn diagram (bottom right) displayed the overlap between genes among FGM32 and MGM4, showing the interesting gene lists in each module. **g**, Dotplot showing the expression of endothelial markers (*VWF* and *PECAM1*) and DE genes comparing F cluster 14 versus F cluster 18 as well as F_GM32 gene module score in F cluster 14 and 18 of Dong_2020_1. The redder the dot, the higher the expression/module score. The size of dots reflected the fraction of cells in corresponding clusters. The asterisk indicated the significance with *p* value < 0.05.

Strikingly, angiogenesis FGMs—notably FGM32—also clustered with immune-response FGMs (Fig. 4b), suggesting functional crosstalk. Indeed, MGM4, a pan-immune module, contained 40% non-immune FGMs (Extended Data Fig. 5c, d), including FGM32, which shared little overlapping genes with its parent MGM (Fig. 4f). This hierarchical divergence implicated FGM32 as an immune-activated angiogenesis signature, independent of classical endothelial markers (*VWF, PECAM1*).

Validation in the Dong_2020_1 dataset revealed two endothelial subpopulations (F clusters 14 vs. 18) distinguished by FGM32 activity (Extended Data Fig. 6b, c). F cluster 14 uniquely upregulated MHC class II genes (*HLA-DRB1, HLA-DPB1*), *CD74*, and *IL3RA* (Fig. 4g; Extended Data Fig. 6c), hallmarks of antigen-presenting endothelial observed in inflammatory niches^30,31^. The strong correlation between MGM4 (immune) and FGM32 (angiogenesis) activity (Fig. 4f, bottom left) further supports their functional synergy. This hierarchical resolution thus uncovers rare, immune-competent endothelial cells—a population obscured in traditional analyses and less mentioned by previous studies^32,33^—demonstrating how SuperSCC bridges broad modules to context-specific states, exposing latent biology in atlas-scale data.

## Discussion

The rapid expansion of single-cell RNA sequencing (scRNA-seq) datasets demands methods that not only scale computationally but also extract biologically meaningful insights from increasingly complex data. SuperSCC addresses this challenge by unifying hierarchical cell-state mapping, conserved gene module (GM) discovery, and automated biological interpretation into a single framework. Crucially, our integration-free approach avoids the pitfalls of batch correction while enabling atlas-scale analyses that reveal cross-tissue biological programs and novel cellular states.

SuperSCC’s hierarchical structure also enables deep exploration of immune contexts, uncovering immune activation even in non-immune cells. Applying SuperSCC to 99 datasets across 20 tissues revealed conserved GMs that define fundamental cellular processes. Immune response and angiogenesis emerged as pan-tissue MGMs, present in >70% of datasets, while finer-resolution FGMs uncovered novel stress-response (e.g., heat shock genes *HSPA1A, HSPB1*) and metabolic programs in immune cells. Strikingly, hierarchical analysis linked immune response MGMs to non-immune FGMs, such as angiogenesis (FGM32) and antioxidant pathways (FGM54/55), revealing cross-compartment coordination. These findings illustrate how SuperSCC’s multi-level framework deciphers cellular plasticity, capturing dynamic transitions and microenvironmental crosstalk.

Notably, SuperSCC identified GMs with direct clinical relevance. Stress-response FGMs conserved across inflammatory diseases correlated with poor immunotherapy response in independent cohorts, while immune-exhaustion modules in epithelial cells aligned with fibrotic progression. Such conserved programs, validated across >50 datasets, highlight SuperSCC’s ability to distill actionable biological themes from noise-rich single-cell data.

We also pioneered the use of large language models (LLMs) to evaluate the gene modules identified by SuperSCC and benchmark them against other methods. The LLM facilitated the validation of SuperSCC-derived modules, improving the accuracy and biological relevance of the results. In the future, like the recent pioneer study^34^, we envision integrating more bioinformatics into an agent-based workflow, where text outputs such as gene names are sent to the LLM for further analysis. While SuperSCC offers significant advances, further improvements are possible. Integrating spatial transcriptomics could provide a more detailed understanding of gene activity in tissue architecture, enabling how gene activity varies across different spatial domains such as tumor margins^35^, immune cell niches^36^, or developmental zones^37^. Besides, incorporating multiomic data types such as genomics, epigenomics, proteomics, and metabolomics—could refine module detection by providing complementary layers of molecular information. For instance, combining DNA methylation data with transcriptomic profiles could uncover regulatory mechanisms driving gene expression changes^38^, while proteomic data could validate the functional outcomes of these changes at the protein level^39^.

Nevertheless, SuperSCC already advances single-cell analysis by bridging hierarchical clustering, cross-dataset GM discovery, and AI-driven interpretation. Its ability to expose hidden biology—from immune-active endothelial to stress-primed immune states—positions it as a critical tool for unraveling complex systems, offering a blueprint for translating ever-growing atlas-scale data into mechanistic insights.

## Acknowledgement

This work was supported by the National Key R&D Program of China (grant no.2023YFF1204701) as well as Major Project of Guangzhou National Laboratory (grant no. GZNL2024A01015 and GZNL2024A03001). We acknowledge the support of the Data Science Platform of Guangzhou National Laboratory and the Bio-medical Big Data Operating System (Bio-OS)^40^.

## Data availability

All data used in this manuscript could be publicly assessed by the URLs stored in Supplementary Table 1.

## Code availability

SuperSCC is available as an open-source Python package at https://github.com/tf1993614/SuperSCC. Scripts to reproduce figures are also available at https://github.com/tf1993614/SuperSCC/tree/main/paper.

## Figures and legends

**Extended Data Fig 1.**
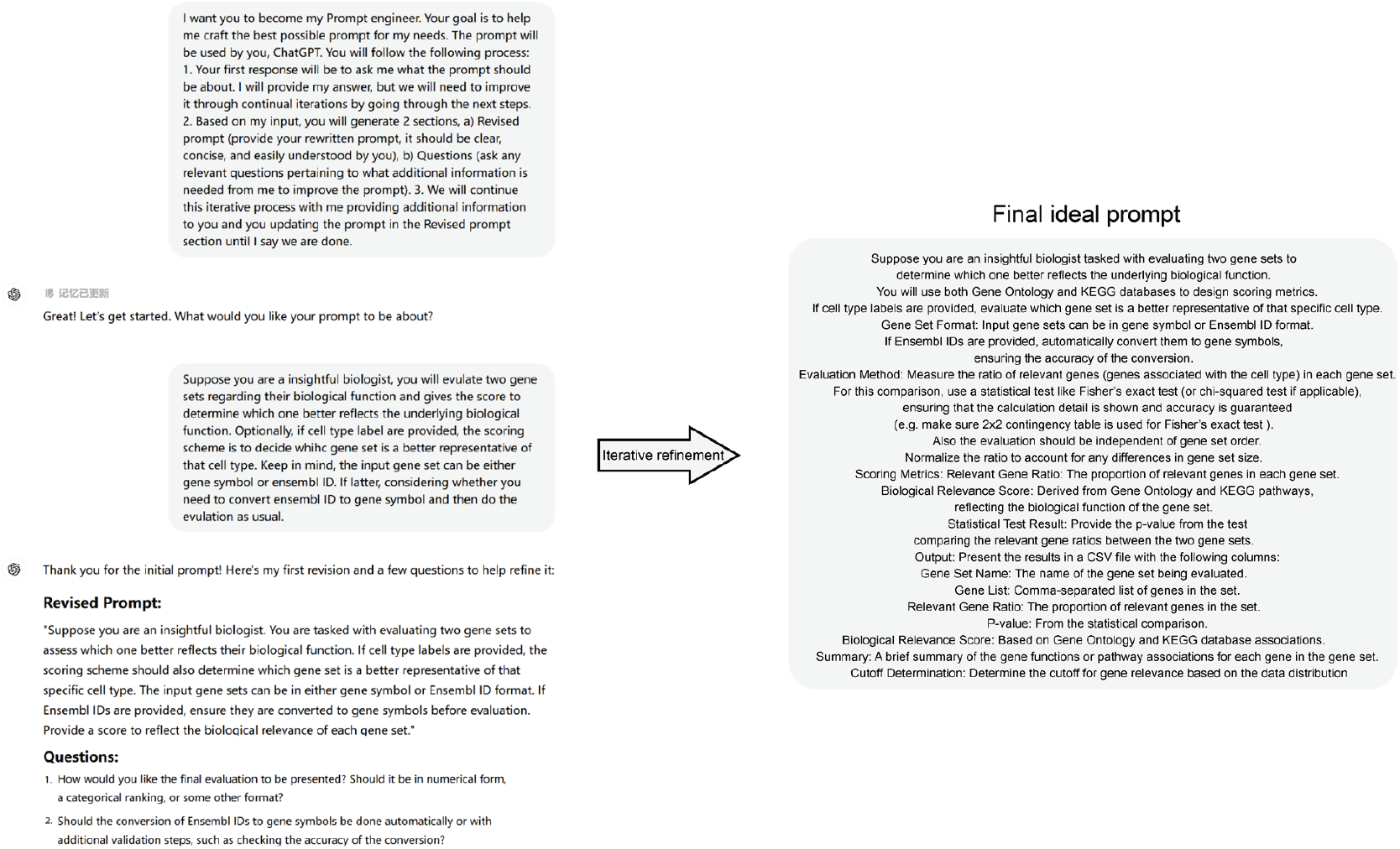
Overview of prompt engineering for obtaining ideal prompt of evaluating gene sets. With making LLM as prompt engineer, we input the first prompt. Then, the LLM would give the revised prompt with improving clarity, conciseness and sustainability by itself. Besides, a few relevant questions would be provided for what additional information from the user to improve the prompt, followed by updating the prompt. Such an iterative refinement process continued until an ideal prompt appeared.

**Extended Data Fig 2.**
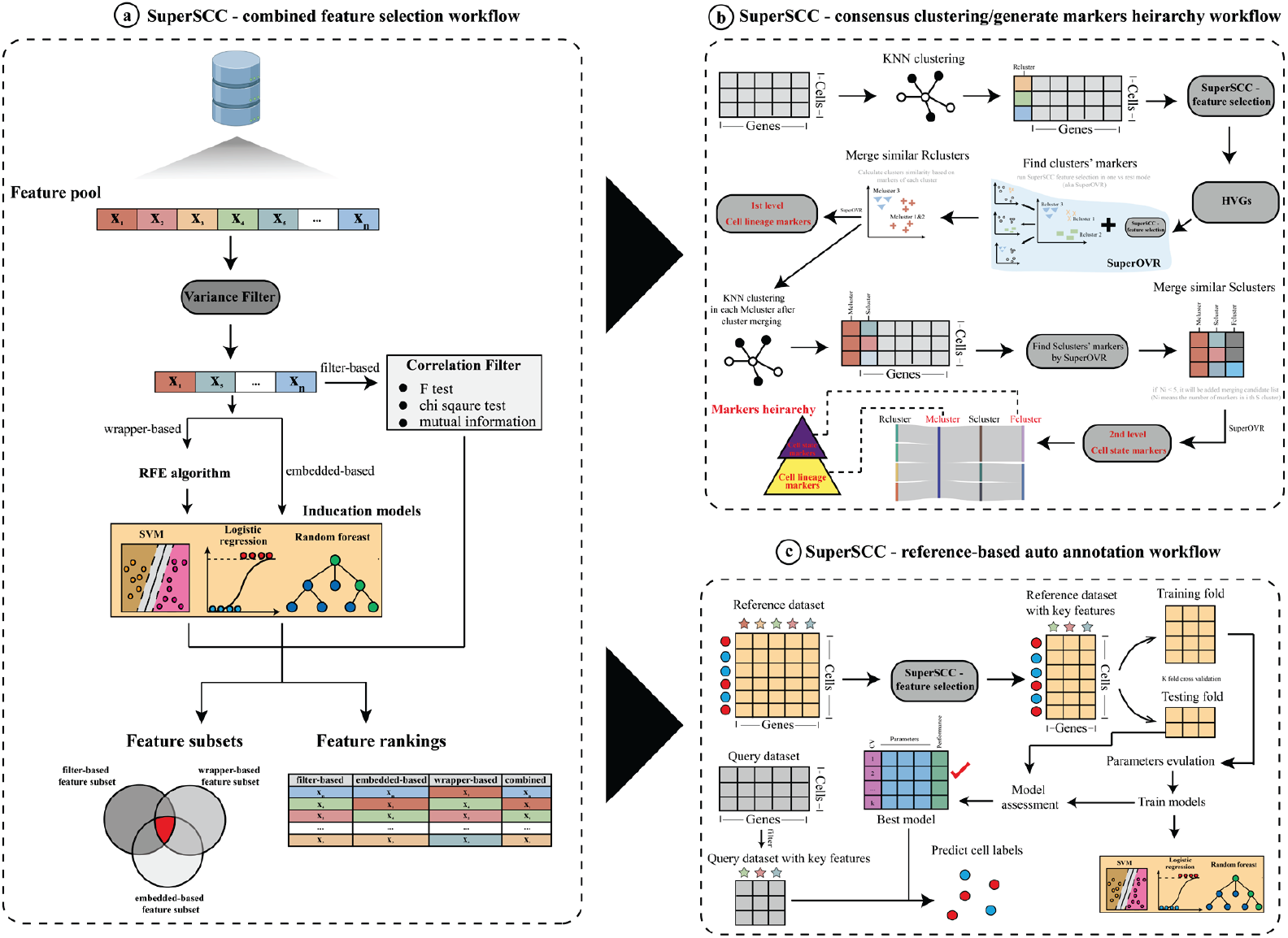
The schematic overview of SuperSCC framework. a, Combined feature selection workflow. Raw features processed with variance filter to remove features with little variability, followed by filter-, embedded- and wrapper-based features selection. With wrapper- and embedded-based categories, three different induction models (SVM, logistic regression and random forest) could be selected. Finally, feature subsets or feature rankings from different feature selection categories were aggregated. **b**, Cluster-aware clustering. R cluster were obtained by graph clustering. Then, with feature selection shown in a, only retaining highly variable genes (HVGs), followed by finding markers in each Rcluster via SuperOVR and further cluster merging thereby forming Mcluster. Such process was repeated in each M cluster to obtain F cluster. With clustering and corresponding markers under broad (M level) and finer (F level) resolution, cell identity and cell markers hierarchy could be established. **c**, Label transfer workflow. Reference dataset processed with features selection in a to remain key features and then split into train fold and testing fold for model parameters evaluation by grid search in cross validation. The best performance model was used to transfer labels in reference to query.

**Extended Data Fig 3.**
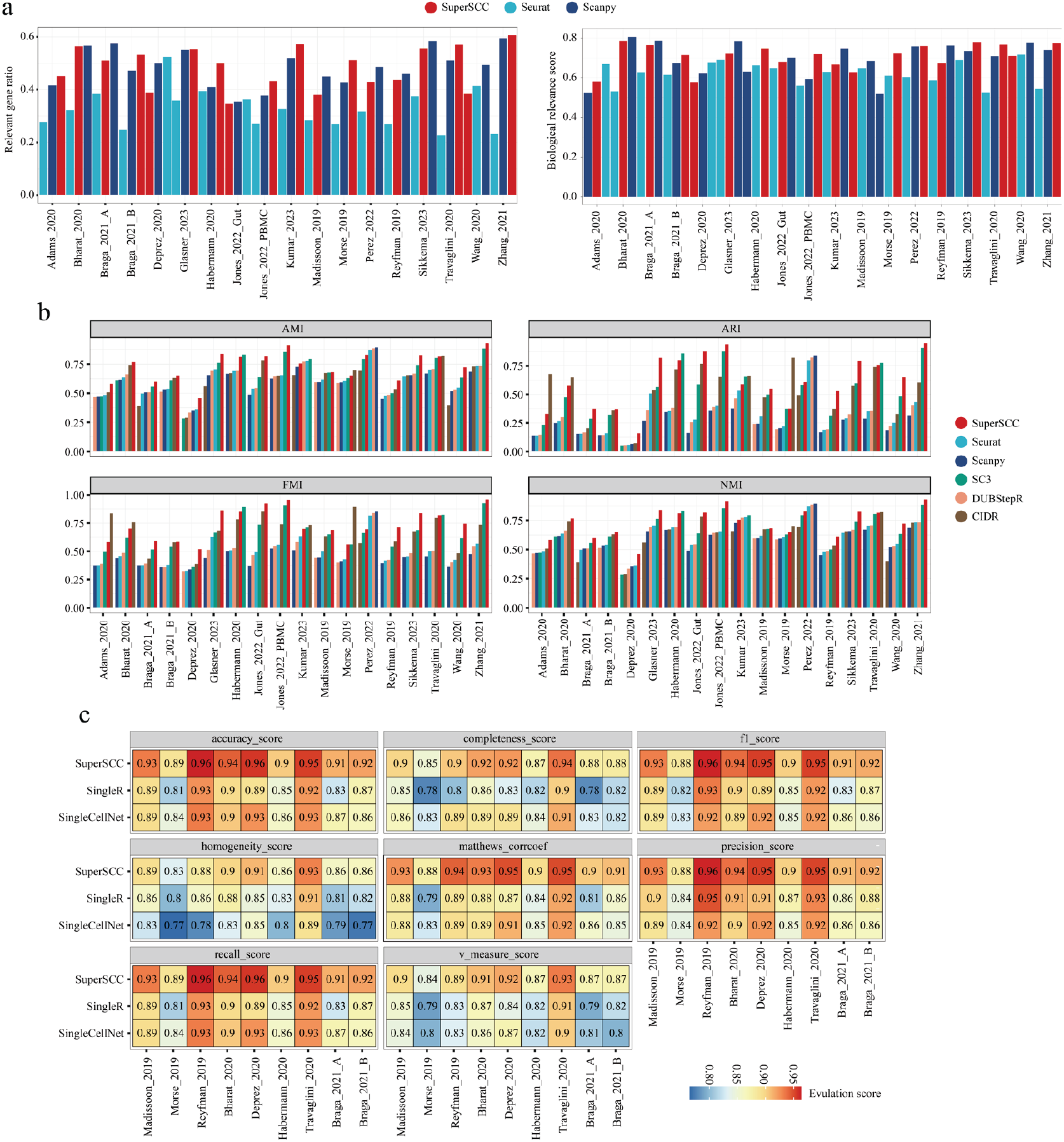
Benchmark SuperSCC with other state-of-art algorithms on different scRNAseq analysis tasks. **a**, Relevant gene ratio (left), biological relevance score (middle) of three comparing algorithms SuperSCC, Scanpy and Seurat on eighteen testing datasets. Bar plots were presented as aggregated average scores on individual testing dataset. **b**, AMI, ARI, FMI and NMI scores of SuperSCC, Scanpy, Seurat, CIDR and DUBStepR on eighteen testing datasets. Bar plots were presented with the aggregated average score on the individual testing dataset. **c**, ACC, Homog, Recall, Prec, Compl, MCC, F1 and VM scores of SuperSCC, SingleR and SingleCellNet on nine testing datasets under self-validation whereby model training and prediction were done on the same dataset. The aggregated average scores were presented per datasets per algorithm. The reader the box, the higher the score.

**Extended Data Fig 4.**
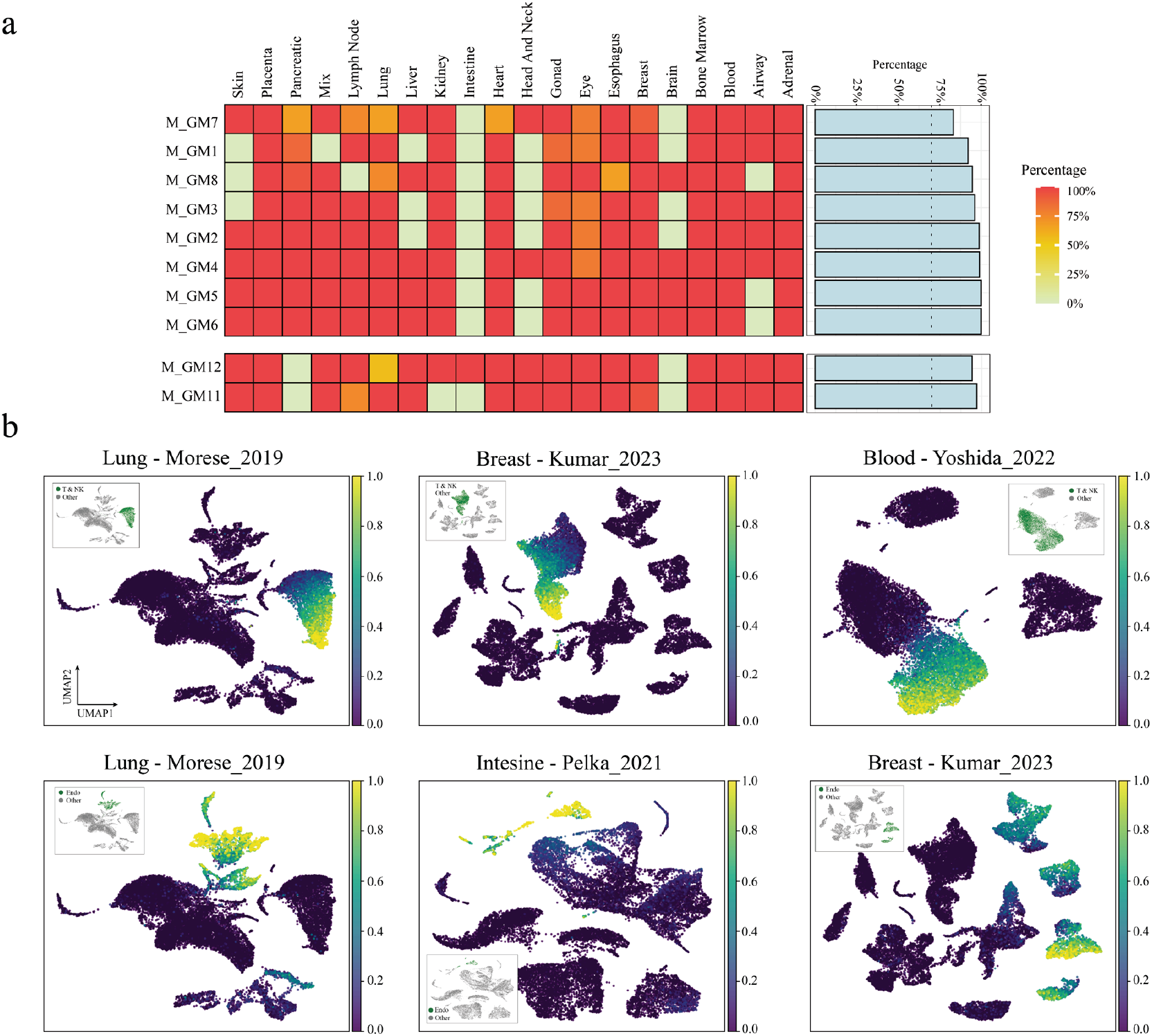
SuperSCC discovers conserved gene modules under M cluster level. **a**, Conservation frequency of each immune response (top) or angiogenesis (bottom) F cluster’s GM (FGM) (rows) in each tissue type (columns). The left annotation bar presented the aggregated average conservation frequency per GM across all tissue types. Color range associated with frequency. The reader the dot, the higher the frequency. **b**, The outer UMAP showing the activity score of immune response MGM6 and angiogenesis MGM12 in a different tissue type in different studies. The more yellow the box, the higher the score. The inner UMAP shows the region of expected cell type (green dots) that were supposed to express the highest activity score of the corresponding FGMs. The cell label was provided by the data generator.

**Extended Data Fig 5.**
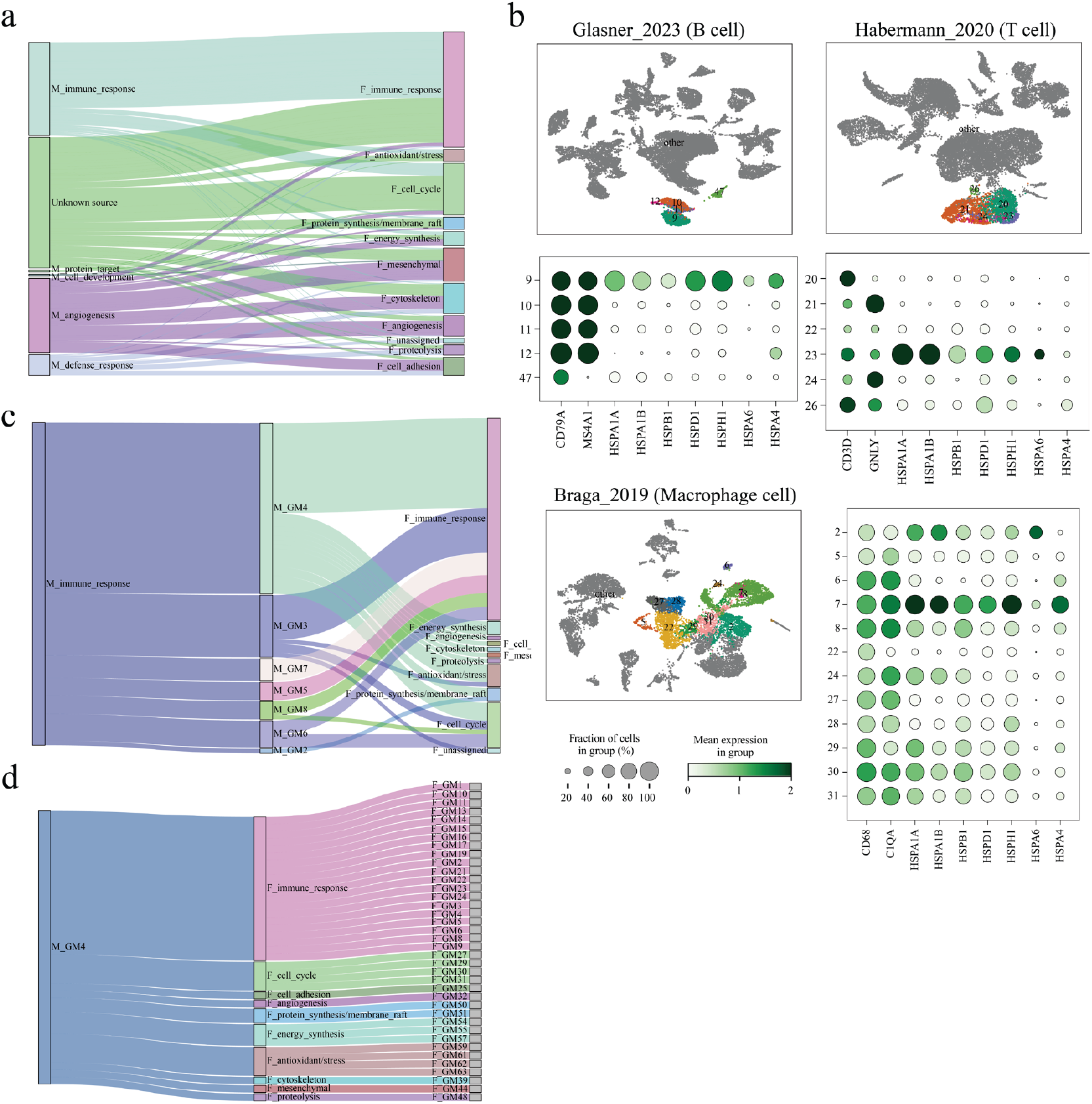
Hierarchical SuperSCC-derived gene modules provide biological insights. **a**, Sankey plot showing the connection between MGMs and FGMs in different functions. **b**, UMAP showing the SuperSCC F clustering with only highlighting the clusters of the focused immune cell types in datasets Glasner_2023 (B cell), Habermanm_2020 (T cell), Brag_2019 (Macrophage cell). Dot plots showing the classical markers of focused immune cell types in each dataset and heat shock genes. The size of dots reflected the fraction of cells in corresponding clusters. The color range represented the average expression of genes in that cluster, whereby the greener the color, the higher the expression. **c**, Sankey plot showing the connection between immune response MGMs and FGMs in different functions. **d**, Sankey plot showing the connection between MGM4 and FGMs in different functions.

**Extended Data Fig 6.**
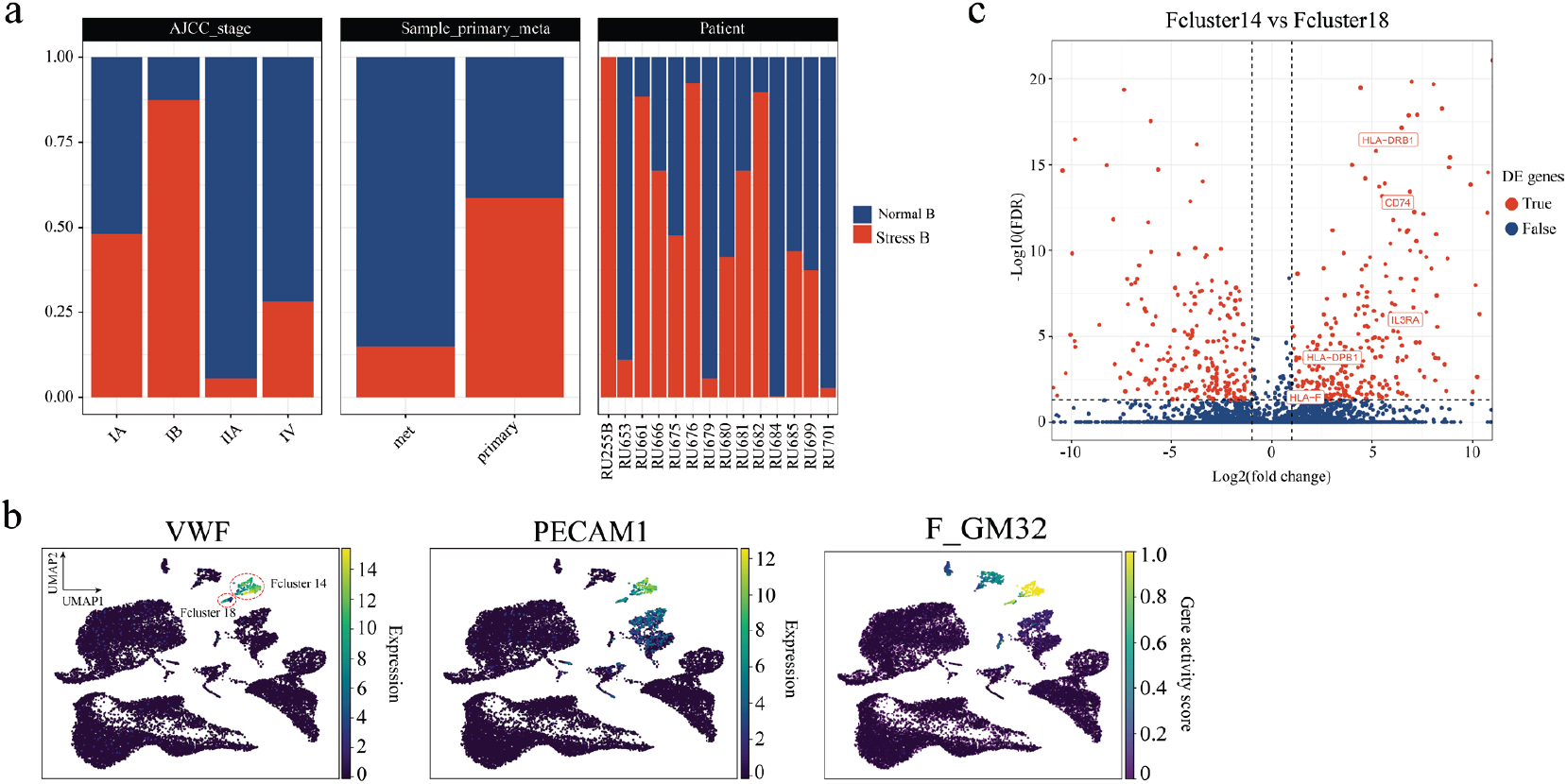
Hierarchical SuperSCC-derived gene modules provide biological insights. **a**, Bar plot showing the frequency of stress B cells under different clinical conditions including tumor stage, tumor metastasis and patient. Color red and blue represented stress and normal B cells respectively. **b**, UMAP showing VWF (top), PECAM1 (middle) genes expression and activity score (bottom) in dataset Dong_2020_1. The dashed line circled SuperSCC F cluster 14 and 18. The yellower the color, the higher the gene expression/gene module activity score. **c**, Volcano plot showing the differential expression genes comparing F cluster 14 versus cluster 18. The red dots denoted the up-regulated genes in F cluster 14 (away from the right vertical dashed line)/Fcluster 18 (away from left vertical dashed line). Interesting up-regulated genes in F cluster 14 were named. The horizontal and vertical dashed line indicated the cutoff for FDR-corrected *p* value (FDR < 0.05) and log2 fold change (|log2foldchange| > 1) respectively.

## Reference

1. Sikkema, L. et al. An integrated cell atlas of the lung in health and disease. Nat Med 29, 1563– 1577 (2023).

2. Tabula Sapiens Consortium* et al. The Tabula Sapiens: A multiple-organ, single-cell transcriptomic atlas of humans. Science 376, eabl4896 (2022).

3. Herring, C. A. et al. Human prefrontal cortex gene regulatory dynamics from gestation to adulthood at single-cell resolution. Cell 185, 4428–4447.e28 (2022).

4. Svensson, V., da Veiga Beltrame, E. & Pachter, L. A curated database reveals trends in single-cell transcriptomics. Database (Oxford) 2020, (2020).

5. Svensson, V., Vento-Tormo, R. & Teichmann, S. A. Exponential scaling of single-cell RNA-seq in the past decade. Nat Protoc 13, 599–604 (2018).

6. Luecken, M. D. et al. Benchmarking atlas-level data integration in single-cell genomics. Nat Methods 19, 41–50 (2022).

7. Argelaguet, R., Cuomo, A. S. E., Stegle, O. & Marioni, J. C. Computational principles and challenges in single-cell data integration. Nat Biotechnol 39, 1202–1215 (2021).

8. Hao, Y. et al. Integrated analysis of multimodal single-cell data. Cell 184, 3573–3587.e29 (2021).

9. Wolf, F. A., Angerer, P. & Theis, F. J. SCANPY: large-scale single-cell gene expression data analysis. Genome Biol 19, 15 (2018).

10. Wang, F., Liang, S., Kumar, T., Navin, N. & Chen, K. SCMarker: Ab initio marker selection for single cell transcriptome profiling. PLoS Comput Biol 15, e1007445 (2019).

11. Kunes, R. Z., Walle, T., Land, M., Nawy, T. & Pe’er, D. Supervised discovery of interpretable gene programs from single-cell data. Nat Biotechnol 42, 1084–1095 (2024).

12. Székely, G. J., Rizzo, M. L. & Bakirov, N. K. Measuring and testing dependence by correlation of distances. Ann. Stat. 35, 2769–2794 (2007).

13. Gavish, A. et al. Hallmarks of transcriptional intratumour heterogeneity across a thousand tumours. Nature 618, 598–606 (2023).

14. Scikit-learn: Machine Learning in Python. Journal of Machine Learning Research 12, 2825–2830 (2011).

15. Kiselev, V. Y. et al. SC3: consensus clustering of single-cell RNA-seq data. Nat Methods 14, 483– 486 (2017).

16. Lin, P., Troup, M. & Ho, J. W. K. CIDR: Ultrafast and accurate clustering through imputation for single-cell RNA-seq data. Genome Biol 18, 59 (2017).

17. Ranjan, B. et al. DUBStepR is a scalable correlation-based feature selection method for accurately clustering single-cell data. Nat Commun 12, 5849 (2021).

18. Aran, D. et al. Reference-based analysis of lung single-cell sequencing reveals a transitional profibrotic macrophage. Nat Immunol 20, 163–172 (2019).

19. Tan, Y. & Cahan, P. SingleCellNet: A Computational Tool to Classify Single Cell RNA-Seq Data Across Platforms and Across Species. Cell Syst 9, 207–213.e2 (2019).

20. DeepSeek-AI et al. DeepSeek-V3 Technical Report. arXiv [cs.CL] Preprint at http://arxiv.org/abs/2412.19437 (2024).

21. Wu, T. et al. clusterProfiler 4.0: A universal enrichment tool for interpreting omics data. Innovation (Camb) 2, 100141 (2021).

22. Kruchten, N., Seier, A. & Chris, P. An interactive, open-source, and browser-based graphing library for Python. https://github.com/plotly/plotly.py doi:10.5281/zenodo.14503524.

23. Wickham, H. ggplot2: Elegant Graphics for Data Analysis. (Springer-Verlag New York, 2016).

24. Veldt, N., Gleich, D. F. & Wirth, A. Learning resolution parameters for graph clustering. arXiv [cs.SI] (2019).

25. DeTomaso, D. & Yosef, N. Hotspot identifies informative gene modules across modalities of single-cell genomics. Cell Syst 12, 446–456.e9 (2021).

26. Chu, Y. et al. Pan-cancer T cell atlas links a cellular stress response state to immunotherapy resistance. Nat Med 29, 1550–1562 (2023).

27. Long, S. et al. Heat Shock Protein Beta 1 is a Prognostic Biomarker and Correlated with Immune Infiltrates in Hepatocellular Carcinoma. Int J Gen Med 14, 5483–5492 (2021).

28. Scutigliani, E. M., Lobo-Cerna, F., Mingo Barba, S., Scheidegger, S. & Krawczyk, P. M. The Effects of Heat Stress on the Transcriptome of Human Cancer Cells: A Meta-Analysis. Cancers (Basel) 15, (2022).

29. Dong, Y. et al. HSPA1A, HSPA2, and HSPA8 Are Potential Molecular Biomarkers for Prognosis among HSP70 Family in Alzheimer’s Disease. Dis Markers 2022, 9480398 (2022).

30. Collins, T. et al. Immune interferon activates multiple class II major histocompatibility complex genes and the associated invariant chain gene in human endothelial cells and dermal fibroblasts. Proc Natl Acad Sci U S A 81, 4917–4921 (1984).

31. Rouhani, S. J. et al. Roles of lymphatic endothelial cells expressing peripheral tissue antigens in CD4 T-cell tolerance induction. Nat Commun 6, 6771 (2015).

32. Xu, Z.-J. et al. Construction of S100 family members prognosis prediction model and analysis of immune microenvironment landscape at single-cell level in pancreatic adenocarcinoma: a tumor marker prognostic study. Int J Surg 110, 3591–3605 (2024).

33. Hamada, A., Torre, C., Drancourt, M. & Ghigo, E. Trained Immunity Carried by Non-immune Cells. Front Microbiol 9, 3225 (2018).

34. Schaefer, M. et al. Multimodal learning of transcriptomes and text enables interactive single-cell RNA-seq data exploration with natural-language chats. bioRxiv (2024) doi:10.1101/2024.10.15.618501.

35. Sibai, M. et al. The spatial landscape of cancer hallmarks reveals patterns of tumor ecological dynamics and drug sensitivity. Cell Rep 44, 115229 (2025).

36. Sun, P. et al. STMiner: Gene-centric spatial transcriptomics for deciphering tumor tissues. Cell Genom 5, 100771 (2025).

37. Zhang, Y. et al. SpaTopic: A statistical learning framework for exploring tumor spatial architecture from spatially resolved transcriptomic data. Sci Adv 10, eadp4942 (2024).

38. Migliaccio, G. et al. Methylation and transcriptomic profiling reveals short term and long term regulatory responses in polarized macrophages. Comput Struct Biotechnol J 25, 143–152 (2024).

39. Jorssen, J. et al. Single-cell proteomics and transcriptomics capture eosinophil development and identify the role of IL-5 in their lineage transit amplification. Immunity 57, 1549–1566.e8 (2024).

40. Liu, J. et al. Bio-medical big data Operating System (bio-OS): An integrated data mining environment for data intensive scientific research. bioRxiv (2024) doi:10.1101/2024.10.17.612997

